# From Homogeneity to Heterogeneity: Refining Stochastic Simulations of Gene Regulation

**DOI:** 10.1101/2024.09.24.614828

**Authors:** Seok Joo Chae, Seolah Shin, Seunggyu Lee, Jae Kyoung Kim

## Abstract

Cellular processes are intricately controlled through gene regulation, which is significantly influenced by intrinsic noise due to the small number of molecules involved. The Gillespie algorithm, a widely used stochastic simulation method, is pervasively employed to model these systems. However, this algorithm typically assumes that DNA is homogeneously distributed throughout the nucleus, which is not realistic. In this study, we evaluated whether stochastic simulations based on the assumption of spatial homogeneity can accurately capture the dynamics of gene regulation. Our findings indicate that when transcription factors diffuse slowly, these simulations fail to accurately capture gene expression, highlighting the necessity to account for spatial heterogeneity. However, incorporating spatial heterogeneity considerably increases computational time. To address this, we explored various stochastic quasi-steady-state approximations (QSSAs) that simplify the model and reduce simulation time. While both the stochastic total quasi-steady state approximation (stQSSA) and the stochastic low-state quasi-steady-state approximation (slQSSA) reduced simulation time, only the slQSSA provided an accurate model reduction. Our study underscores the importance of utilizing appropriate methods for efficient and accurate stochastic simulations of gene regulatory dynamics, especially when incorporating spatial heterogeneity.

## 1. Introduction

Gene regulation is a fundamental process that governs cellular functions, organismal development, and responses to environmental changes [1, 2, 3]. In eukaryotes, this process is primarily regulated by interactions between DNA and DNA-binding proteins (e.g., transcription factors) within the nucleus [4, 5, 6, 7]. Due to the limited number of molecules involved in the nucleus, often including just one or two copies of DNA and a few dozen DNA-binding proteins, the stochastic nature of gene regulation becomes significant [8]. This stochasticity leads to fluctuations in gene expression, which can be crucial for processes such as cell division and environmental adaptation. To comprehend the impact of stochasticity on gene regulation, the chemical master equation (CME), which describes the probabilities of different molecular states over time, has been frequently used. However, analytical solutions to the CME are typically challenging to obtain [9, 10, 11, 12, 13, 14, 15]. To address this, the stochastic simulation algorithm (SSA) [16], represented by the Gillespie algorithm, has been widely used for simulating gene regulation [17, 18, 19, 20, 21, 22, 23, 24, 25, 26, 27, 28].

The SSA assumes homogeneity within the nucleus, treating molecules as evenly distributed. However, in reality, genes are localized at specific locations within the nucleus rather than being uniformly distributed [29, 30]. This spatial heterogeneity may affect the accuracy of the SSA in describing actual transcription processes. The impact of spatial heterogeneity is expected to be minimal when diffusion of proteins occurs rapidly. However, under slow diffusion conditions, assuming a homogeneous distribution can lead to significant deviations from reality.

When the SSA does not accurately capture the effects of spatial heterogeneity, it is crucial to employ alternative stochastic simulation methods that incorporate spatial heterogeneity. Previous studies have addressed spatial heterogeneity using either compartment-based or agent-based methods. Compartment-based methods, such as compartment-based SSA [31], URDME [32], MesoRD [33], SmartCell [34], and Lattice Microbes [35], divide the domain into small compartments where diffusion and reactions are modeled using the SSA or its variants. On the other hand, agent-based methods like STEPS [36] and Smoldyn [37] simulate the Brownian motion of individual molecules and reactions that occur upon molecular collisions. By incorporating spatial heterogeneity, these methods have effectively described complex biological systems.

Despite accurate descriptions of systems, stochastic simulations become computationally intensive when spatial heterogeneity is incorporated. To reduce this computational burden, the Langevin approach, which simplifies the SSA, has recently been proposed for application to spatial SSA [38, 39, 40]. However, this method may not be suitable for systems describing gene regulation as it typically requires a large number of molecules. In such cases, the quasi-steady-state approximation (QSSA) presents a promising alternative. Various QSSA methods have successfully accelerated the SSA by simplifying gene regulation models [9, 10, 11, 12, 13, 14, 41, 42, 43, 44, 45, 46, 47, 48, 49, 50, 51]. However, it remains unclear whether QSSA can be effectively applied to the spatial SSA.

In this study, we evaluated the accuracy of the SSA in describing gene regulation within heterogeneous environments by comparing the standard SSA with the spatial SSA. To simulate the spatial SSA, we employed the compartment-based SSA [31], since the agent-based SSA can be infeasible for a large number of molecules, whereas the compartment-based approach remains feasible [52, 53]. Our findings indicate that the SSA and the spatial SSA produce different outcomes when transcription factors diffuse slowly. Furthermore, when these transcription factors are non-uniformly distributed in the nucleus or when there are multiple DNA binding sites, this difference becomes larger. Additionally, we examined whether model reduction techniques for SSA can be extended to the spatial SSA. We found that using the stochastic total quasi-steady-state approximation (stQSSA) to reduce the gene regulation model is accurate for the SSA but yields significant errors for the spatial SSA. On the other hand, the reduced model using the stochastic low-state quasi-steady-state approximation (slQSSA) provides accurate simulations for both the SSA and the spatial SSA, particularly when gene copy numbers are small. Our study provides guidelines on how to simulate and simplify models describing gene regulation by transcription factors with slow diffusion in spatially heterogeneous environments.

## 2. Results

### 2.1. Gene regulation models can be simulated with SSA or spatial SSA

Transcriptional regulation is a key mechanism of gene expression, often involving transcription factors binding to DNA [3]. These factors can either promote or inhibit transcription. Our study focuses on transcriptional repressors within a simple gene regulatory network. To illustrate this, we used an established model where a transcription factor inhibits gene expression [27].

As a basic case, we modeled gene regulation of DNA with a single binding site (Fig. 1A). In this model, the DNA (*D*_0_) has one binding site that can reversibly bind to a repressor (*P*) with a binding rate of *k*_*f*_ and an unbinding rate of *k*_*b*_. When the binding site is occupied (*D*_1_), transcription is inhibited, but when the site is unoccupied, transcription proceeds at a rate of *k*_*p*_. The produced mRNA subsequently degrades at a rate of *k*_*d*_.

**Figure 1.**
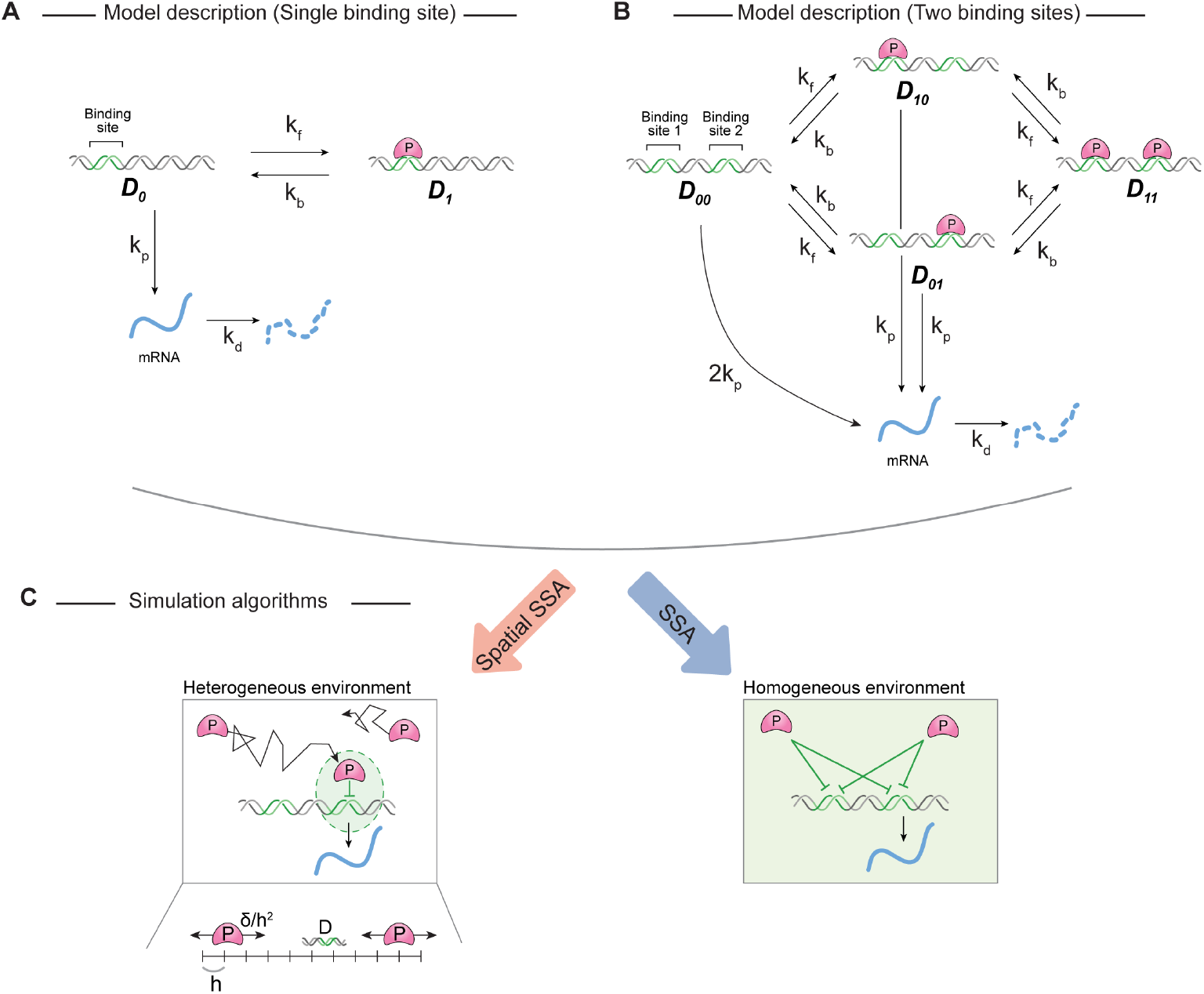
Frameworks for stochastic simulation of gene regulation with and without spatial heterogeneity. **(A)** Schematic diagram of the model describing the repressor (*P*) to a binding site on DNA. *P* can reversibly bind to an unoccupied site on the DNA (*D*_0_) with a binding rate *k*_*f*_ and an unbinding rate *k*_*b*_. Unoccupied DNA (*D*_0_) is transcribed at a rate of *k*_*p*_. When the binding site is occupied (*D*_1_), the transcription is repressed. Transcribed mRNA decays at the rate of *k*_*d*_. **(B)** When the DNA has two binding sites, *P* can reversibly bind to either site. If both binding sites are unoccupied (*D*_00_), mRNA is transcribed with the rate of 2*k*_*p*_. If one site is occupied (*D*_01_ or *D*_10_), the transcription rate is halved. *P* can still reversibly bind to the other site with the same binding rates (*k*_*f*_), indicating that the binding of *P* to each site occurs independently. When both sites are occupied (*D*_11_), transcription is fully repressed. **(C)** These models can be simulated using either the spatial SSA (left) or the SSA (right). The spatial SSA describes not only the reactions but also the diffusion of *P* (left). To describe diffusion, we divided the domain into multiple compartments of size *h*. The diffusion rate is calculated by *δ/h*^2^, where *δ* is the diffusion coefficient of *P*. *P* is required to diffuse to DNA before binding, ensuring that only *P* in proximity to DNA binding sites can bind. In contrast, the SSA assumes a homogeneous environment for DNA and *P*, allowing *P* to bind to DNA binding sites from anywhere in the nucleus (right).

In gene regulation, it is common for DNA to contain multiple binding sites for transcription factors [54, 55, 56]. To describe gene regulation involving multiple binding sites, we first examined a simple model with DNA containing two binding sites (Fig. 1B and Eq. 1). When the DNA is unoccupied (*D*_00_), transcription occurs at a rate of 2*k*_*p*_. If either binding site is occupied (*D*_01_ or *D*_10_), the transcription rate is halved. *P* can bind to the remaining site, and when both binding sites are occupied (*D*_11_), transcription is fully suppressed. Since the model distinguishes all DNA binding statuses, simulation and analysis can be complex. Given this complexity, we sought a way to simplify the model while maintaining its essential dynamics. To achieve this, we leveraged the fact that the transcription rate is proportional to the number of free binding sites. For example, mRNA is transcribed from *D*_00_, which contains two free binding sites, at a rate of 2*k*_*p*_. Similarly, mRNA is transcribed at a rate of *k*_*p*_ from *D*_01_ and *D*_10_. Thus, by introducing *D*, which is the number of unoccupied binding sites, we can describe the transcription from various statuses of DNA (*D*_00_, *D*_01_, *D*_10_, and *D*_11_) by simply using *k*_*p*_*D*. That is, we can describe numerous transcription reactions from various DNA binding statuses with a single reaction from *D*. This approach allows us to construct a simpler model (Eq. 4) with dynamics identical to the original (Eq. 1) (see Methods for details).

To simulate gene regulation models, two methods can be utilized: the spatial SSA and the SSA (Fig. 1C). The spatial SSA, based on the compartment-based Gillespie algorithm [31], describes the diffusion and reactions of species. Specifically, the spatial SSA divides the domain into compartments of size *h*, and *P* can diffuse across compartments at a rate of *δ/h*^2^, where *δ* is the diffusion coefficient of *P*. The binding of *D* and *P* occurs only when both are in the same compartment. While the spatial SSA provides a detailed simulation, it can be computationally intensive. To circumvent this, previous studies employed the Gillespie algorithm to implement the SSA for computational efficiency. The SSA assumes a homogeneous environment where all *P* can bind to the sites with the same propensity, ignoring spatial heterogeneity. While this simplifies the simulation and speeds it up, using the SSA may capture the gene regulation inaccurately when spatial heterogeneity plays a role.

### 2.2. Utilizing SSA results in an error in simulating gene regulation models when the diffusion of transcription factor is slow

While the SSA assumes that a nucleus is a well-mixed environment, it is a biologically irrelevant assumption since DNA is localized in a specific location. To investigate whether the SSA can still accurately capture transcription, we compared the SSA with the spatial SSA. By simulating the gene regulation model (Fig. 1) by the SSA and the spatial SSA, we quantitatively compared mRNA levels over time under various initial conditions. Specifically, we compared the mean and standard deviation of mRNA amounts across 1,000 simulation iterations for each method.

We first simulated from a simple model where *D* is localized in the center of the nucleus and *P* is uniformly distributed across the nucleus initially (Fig. 2A). When diffusion is fast, the SSA and the spatial SSA produce similar means and standard deviations of mRNA amounts (Fig. 2B). This situation resembles well-mixed conditions, resulting in no significant differences between the SSA and the spatial SSA.

**Figure 2.**
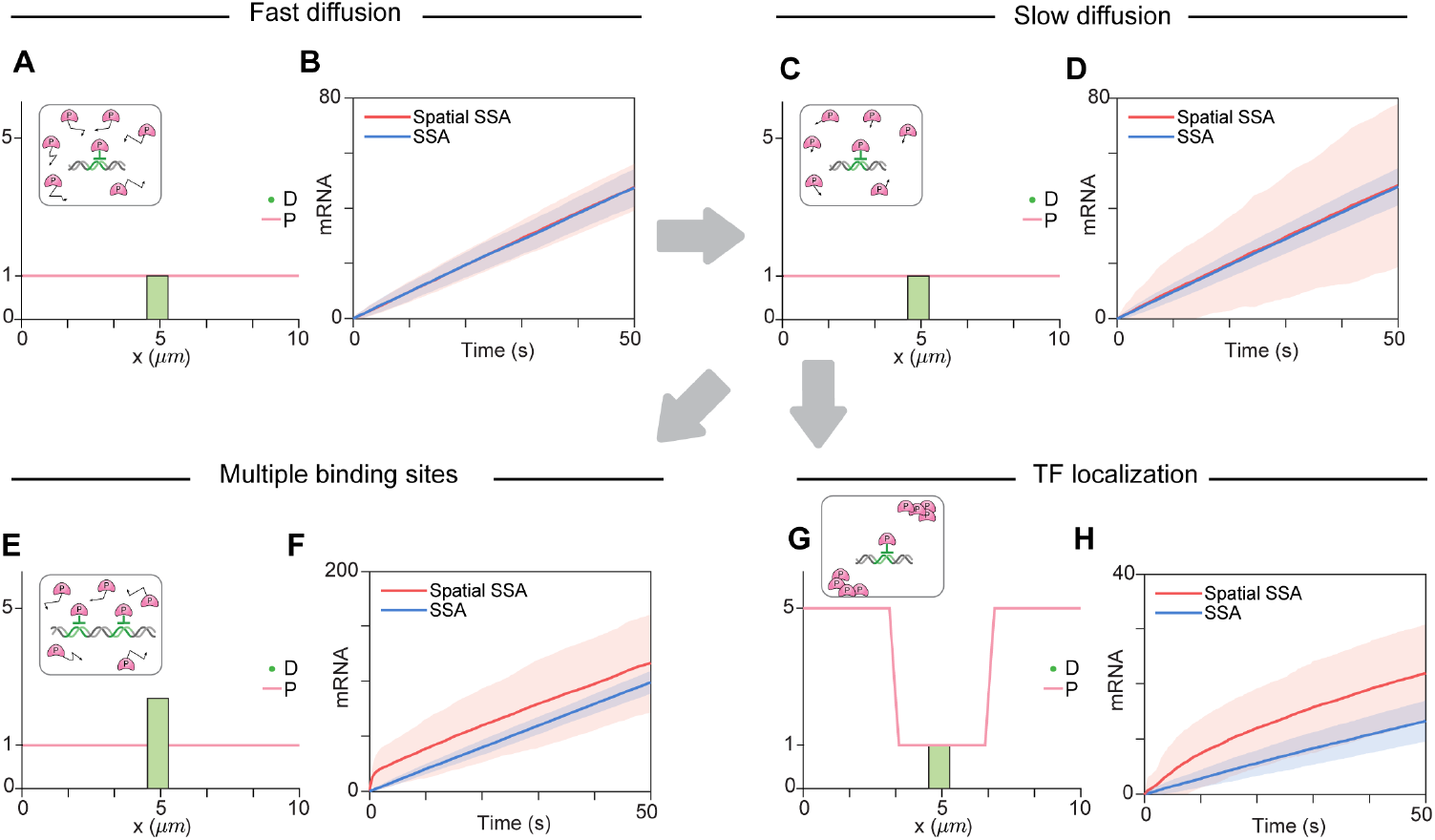
Utilizing the spatial SSA is essential for accurately simulating gene regulation with multiple binding sites or the non-uniform distribution of transcription factors within the nucleus. **(A)** The regulation of DNA with a single binding site, when *P* is uniformly distributed, is described by the initial condition where *P* = 1 throughout the nucleus and *D* = 1 at the nucleus center. The diffusion coefficient of 10*µm*^2^*/s* lies in the typical range of protein diffusion coefficients. **(B)** With these initial conditions, both the spatial SSA and the SSA result in the same average mRNA production and the same standard deviation of mRNA production over time. This occurs because when *P* is uniformly distributed, *D* and *P* maintain a stoichiometric ratio of 1:1. **(C)** To describe the scenario when diffusion occurs slowly, the same initial conditions as in (A) were used, but the diffusion coefficient was decreased to 0.2*µm*^2^*/s*. **(D)** When diffusion occurs slowly, the SSA results in the same average mRNA production as the spatial SSA; however, the SSA underestimates the standard deviation of the mRNA production. This occurs because diffusion of *P* to *D* results in additional noise in the transcription. **(E)** To describe the regulation of DNA with two binding sites when *P* is uniformly distributed, we utilized the initial condition where *P* = 1 throughout the nucleus and *D* = 2 at the nucleus center. **(F)** With these initial conditions **(E)**, the SSA underestimates the average mRNA production more than the spatial SSA does. This is because the SSA allows transcription factors at every location to bind to the DNA, which results in an overestimation of the number of occupied sites. **(G)** To describe when transcription factors are localized in a specific position, we utilized a non-uniform distribution. Specifically, *P* is assumed to be accumulated (*P* = 5) at the periphery of the nucleus, while *P* = 1 is assumed to be near the nucleus center. **(H)** With these initial conditions, the SSA highly underestimates the mRNA production compared to the spatial SSA.

However, when diffusion becomes slow, the SSA underestimates the standard deviation of the mRNA amount, while it still provides a similar average mRNA production to that of the spatial SSA (Fig. 2C-D). Slow diffusion leads to infrequent transitions between active and inactive DNA states, as *P* moves more slowly in and out of DNA-containing compartments. This results in a longer length of time of either high mRNA production (if the DNA remains active) or no changes in mRNA levels (if the DNA remains inactive). Consequently, the spatial SSA shows larger variations in mRNA amounts compared to the SSA under slow diffusion conditions (Fig. 2D).

However, the average amount of *P* in any compartment containing DNA remains constant since *P* is uniformly distributed, regardless of the diffusion rate. Thus, the average mRNA production is not affected by the diffusion coefficient. Given that the SSA corresponds to a situation where DNA diffuses rapidly across compartments, the spatial SSA and the SSA result in similar average mRNA production. Consequently, when diffusion of *P* is slow and DNA has a single binding site, the SSA predicts the average of the spatial SSA well, but poorly predicts the variance.

We further investigated whether the SSA can accurately describe gene regulation with multiple binding sites. We used an initial condition where two binding sites are located in the center of the nucleus (Fig. 2E). In this case, the SSA underestimates both the average and the variance of mRNA transcription compared to the spatial SSA (Fig. 2F). This discrepancy arises because the SSA does not account for the time it takes for *P* to diffuse and reach the DNA binding sites. When the local concentration of *P* is insufficient to saturate all the binding sites, the SSA inaccurately assumes that every *P* molecule can bind to *D* regardless of its spatial position. However, in reality, only *P* molecules close to *D* can bind. Consequently, the SSA model shows DNA becoming occupied more rapidly than in the spatial SSA model, resulting in greater inhibition. Additionally, when the distribution of *P* is nonuniform, such that the local concentration of *P* near DNA is lower than the average concentration, the SSA underestimates mRNA transcription compared to the spatial SSA (Fig. 2G-H). This underestimation of transcription occurs because the SSA overestimates *P*, thereby overestimating the repression of *D*.

Consequently, using the SSA to describe gene regulation under conditions of slow diffusion, multiple binding sites, and non-uniform distribution of transcription factors can lead to results that differ significantly from those observed in real biological systems.

### 2.3. Stochastic tQSSA can lead to an erroneous reduction of spatiotemporal models

In the previous section, we demonstrated the necessity of the spatial SSA to model gene regulation accurately under slow diffusion. Under these conditions, the majority of computation time is spent simulating fast reactions. Since reversible binding between a transcription factor and DNA typically occurs faster than transcription, these reversible binding reactions become the primary source of computational cost. To address this, we aimed to develop a more efficient method to reduce the spatial SSA by eliminating fast reactions without compromising accuracy.

Accurate methods for eliminating fast reactions in the SSA have been previously reported. These methods are based on the assumption that species involved in rapid binding and unbinding reactions reach their quasi-steady states (QSSs) faster than other species not affected by reversible binding. This assumption allows for replacing the variables involved in fast reactions with their quasi-steady-state approximations (QSSAs), thereby eliminating the fast reactions. However, since these methods are derived for the SSA under the assumption of homogeneous environments, it is unclear whether QSSAs remain accurate in heterogeneous environments. Therefore, in this section, we investigated the applicability of QSSAs derived for the SSA within the spatial SSA, particularly under conditions of slow diffusion.

The QSSAs of species in homogeneous environments are obtained assuming the QSS of binding and unbinding reactions. For stochastic models, the QSSAs for species are obtained as the stationary average number of the species when the system reaches QSS. In our model, we calculate the QSSA of free DNA (*D*_*QSSA*_) as it directly affects the amount of mRNA. Calculating the *D*_*QSSA*_ requires solving the CMEs describing the gene regulation, which is usually highly challenging. However, when the DNA contains few binding sites, it can be solved analytically. This is called the stochastic low-state QSSA (slQSSA) (Fig. 3A) [14]. The slQSSA becomes highly complex and difficult to analyze as the number of binding sites increases. In such cases, the stochastic total QSSA (stQSSA) [10, 11, 12, 13, 14, 42, 44, 45, 46, 47, 48, 49, 50, 51], can be used as an alternative. The stQSSA of *D* is a non-elementary function, which is obtained from the steady-state solution of the associated differential equation in terms of the total variables, *D*_*T*_ = *D* + *D* : *P* [10, 11, 12, 13, 14, 42, 44, 45, 46, 47, 48, 49, 50, 51]. This is then used to calculate propensity functions in the Gillespie algorithm for stochastic simulations. Although the stQSSA is generally accurate in the SSA [10, 11, 12, 13, 14, 42, 44, 45, 46, 47, 48, 49, 50, 51], it has been reported that tight binding with similar amounts of *D* and *P* results in discrepancies between the predicted and actual average free DNA [14]. This error is most pronounced when the ratio of *D* and *P* is 1:1, where the error upper bound peaks (Fig. 3B) [14].

**Figure 3.**
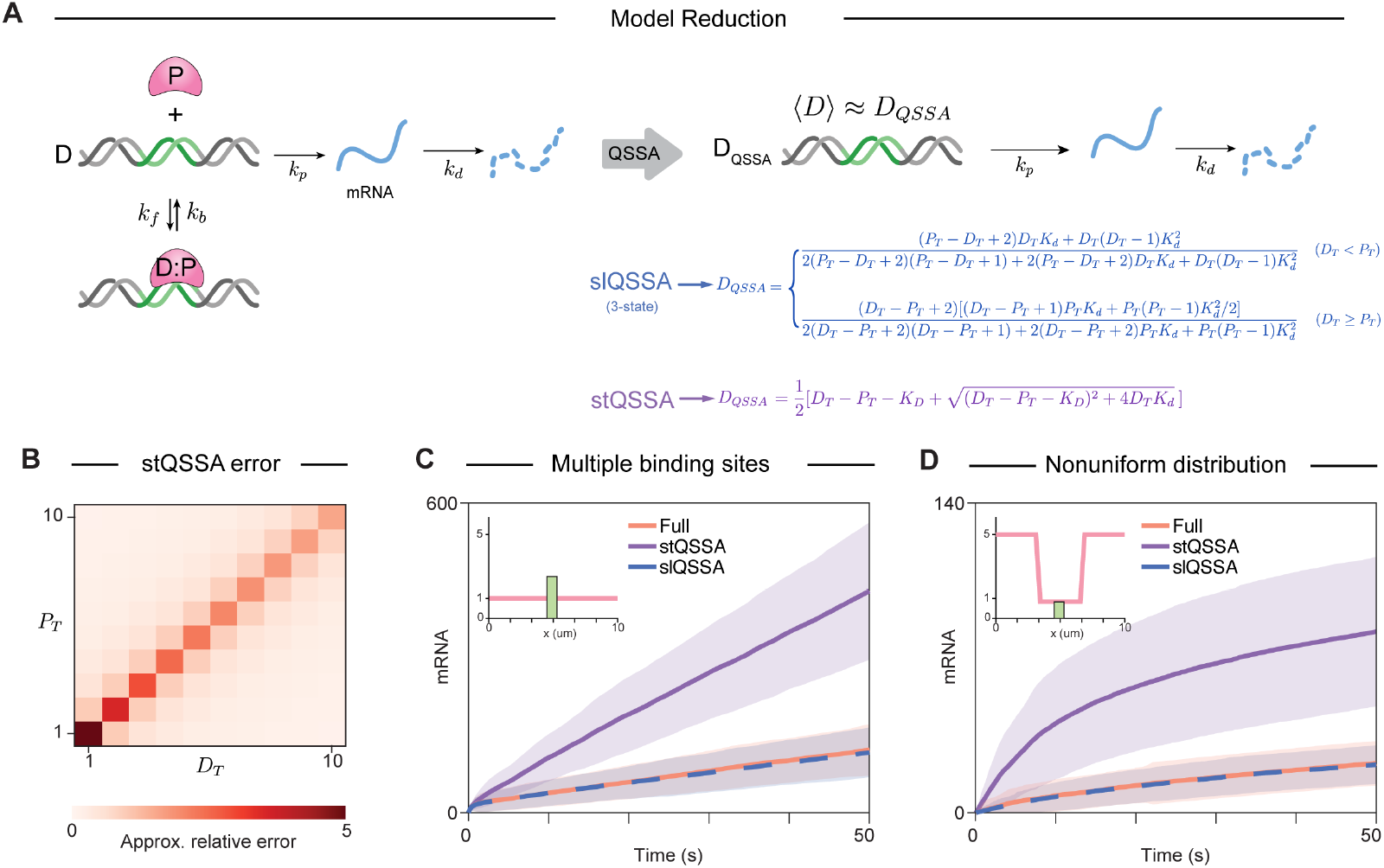
slQSSA, but not stQSSA, provides an accurate reduction of the full model describing gene regulation. **(A)** The full model describing gene regulation, based on mass-action kinetics, can be reduced by replacing *D* by its stQSSA or slQSSA. Both approximations assume that *D* reaches a QSS rapidly, which is determined by the total DNA amount (*D*_*T*_) and the total protein amount (*P*_*T*_). See Methods for details. **(B)** The relative error of the stQSSA increases as the sensitivity of the *D*_*QSSA*_ increases. The stQSSA results in a large sensitivity, and thus a large error when *D*_*T*_*≈P*_*T*_. (C-D) As a result, stQSSA overestimates the average mRNA production in both cases: when a small number of transcription factors interact with the two binding sites (C, inset), and when *D*_*T*_*≈P*_*T*_ due to *P* localization at the nucleus periphery (D, inset). In contrast, slQSSA accurately estimates the average mRNA production within the nucleus, regardless of the initial conditions.

We examined the accuracy of two QSSAs, the slQSSA and the stQSSA, derived for the SSA within the spatial SSA. Specifically, we compared the mRNA levels obtained from a gene regulation model describing multiple binding sites by simulating the full model, the slQSSA model, and the stQSSA model. We chose initial conditions where the overall molar ratio of *D* and *P* deviates from 1:1 (*D* = 2, *P* = 30), a scenario in which the stQSSA is expected to perform well (Fig. 3C, inset).

Unexpectedly, the stQSSA model overestimated the mRNA amount compared to the full model (Fig. 3C). This overestimation occurred because the local ratio of *D* and *P* near the DNA approached 1:1, leading to an overestimation of *D*. In contrast, the slQSSA captured the full model accurately (Fig. 3C). When we repeated the simulation with initial conditions describing a non-uniform distribution of *P*, the stQSSA again resulted in an error (Fig. 3D). This error was due to the local ratio of *D* and *P* approaching 1:1. However, the slQSSA once again accurately captured the full model (Fig. 3D).

Therefore, we found that stQSSA can be inaccurate in the spatial SSA, even under conditions that were predicted to be accurate in the SSA. However, even with the spatial SSA, the slQSSA always correctly reduced the full model, as it is the analytic solution of CMEs. These findings suggest that we should avoid using stQSSA to reduce spatially heterogeneous models.

### 2.4. Stochastic tQSSA can distort oscillatory dynamics

So far, we have shown that the stQSSA does not accurately approximate the full model even when the overall molar ratio of *D* and *P* is different from 1:1 by examining cases when the molar ratio is conserved. However, in real cells, the number of transcription factors changes over time, which can cause the molar ratio to vary. This raises the question of whether the stQSSA is accurate in situations where the molar ratio is 1:1 for only a short period. To investigate this, we examined the accuracy of the stQSSA and the slQSSA in a model exhibiting oscillatory dynamics.

To explore this, we constructed a negative feedback model designed to generate oscillatory behavior (Fig. 4A). Specifically, when *P* is produced, it instantly translocates to the cell periphery and then diffuses to the center of the nucleus, where *D* is located. Upon binding to *D, P* represses its own production, leading to a decrease in its amount. As *P* decreases and *D* becomes unbound, transcription resumes, increasing the amount of *P*. This negative feedback loop induces oscillations through repeated cycles.

**Figure 4.**
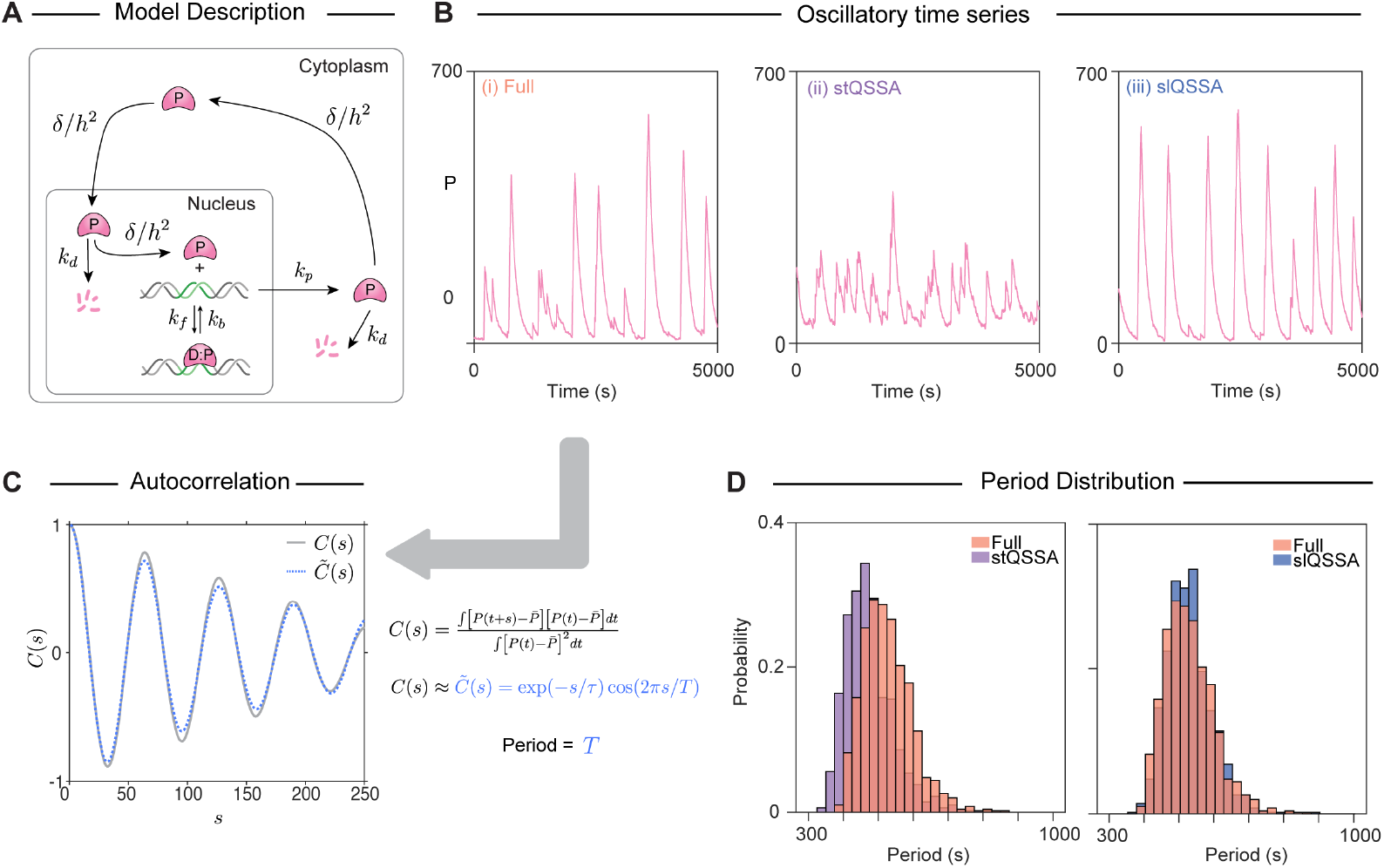
The stQSSA underestimates the period of the biological oscillation. Full model diagram of an oscillatory transcriptional negative feedback loop. From the unoccupied DNA, the repressor (*P*) is produced and then translocated to the periphery of the cytoplasm. The *P* then diffuses to the nucleus. After entering the nucleus, the *P* diffuses to the DNA and binds to it to form a complex, which is transcriptionally inactive, and thus represses its own production. As the reversible binding between *P* and *D* is rapid, by replacing *D* with its QSSAs (either stQSSA or slQSSA), we can obtain reduced models. Oscillatory trajectories of *P* simulated with the full model (i), the stQSSA model (ii), and the slQSSA model (iii). In the stQSSA model, transcription occurs more frequently than in the full model, while the slQSSA model shows transcription occurring at a similar frequency to that of the full model. (C) To measure the period of each time series, the autocorrelation of the time series was calculated. The period was then estimated by fitting the resulting data to a decaying cosine function. (D) The stQSSA model predicts a shorter period than the full model (left). In contrast, the slQSSA model accurately predicts the period of the full model (right). The distribution of periods was obtained from the 1,000 iterations of the simulation.

To evaluate whether the reduced models can accurately capture the oscillations of the full model, we simulated these models using the spatial SSA (Fig. 4A). The oscillations of *P* in the full model (Fig. 4B(i)) are not accurately captured by the stQSSA (Fig. 4B(ii)). The inaccuracy of the stQSSA model arises because transcription occurs more frequently when the molar ratio is close to 1:1 (Fig. 3C-D), leading to faster repression and a shorter oscillation period. In contrast, the slQSSA model closely captures the oscillations of the full model (Fig. 4B(iii)).

To quantify the period of the time series obtained from each model, we utilized the autocorrelation of each time series [28, 57, 58]. Specifically, we estimated the period by fitting the oscillatory time series to a decaying cosine function (Fig. 4C). The stQSSA model underestimated the oscillation period of the full model, while the slQSSA model accurately predicted the oscillation period (Fig. 4D). This indicates that the stQSSA can distort oscillatory dynamics even if the 1:1 ratio is maintained for only a short time. On the other hand, the slQSSA remains accurate, making it a more reliable choice for properly reducing spatially heterogeneous models.

## 3. Discussion

Stochasticity plays a crucial role in gene regulation due to the limited number of molecules involved in the process [3]. To understand this phenomenon, previous studies have extensively employed the SSA, which assumes the homogeneity of the system [17, 18, 19, 20, 21, 22, 23, 24, 25, 26, 27, 28]. Our study investigated whether this assumption is appropriate for accurately describing biological systems. Our findings reveal that when transcription factors diffuse slowly, the SSA fails to adequately capture spatial heterogeneity within the cellular environment (Fig. 2C-D). Conversely, when the diffusion of transcription factors is rapid compared to reaction rates, the spatial SSA and the SSA yield similar results (Fig. 2A-B). This is because when the diffusion timescale is shorter than the reaction timescale, the system becomes homogenized before reactions affect the system appreciably. Typically, reaction timescales for binding, unbinding, and catalysis are under 1 *s*. Given that the size of a eukaryotic nucleus (*L*) is about 10 *µm* [59], and the diffusion coefficient of transcription factors (*D*) ranges from 0.5 to 5 *µm*^2^*/s* [59, 60], the diffusion time scale (= *L*^2^*/D*) is estimated to be approximately 20 to 200 *s*. Therefore, the diffusion time scale of transcription factors is generally slower than the reaction time scale. Thus, using the spatial SSA is crucial for accurately simulating gene regulation.

Stochastic simulations often require significantly more time than deterministic methods, necessitating acceleration techniques. Various QSSA methods, such as the sQSSA, stQSSA, and slQSSA, have been employed to speed up simulations in the SSA [9, 10, 11, 12, 13, 14, 41, 42, 43, 44, 45, 46, 47, 48, 49, 50, 51]; however, their effectiveness in the spatial SSA has been elusive. Given the success of tQSSA in deterministic partial differential equation simulations [61, 62], it seemed a natural candidate for application to stochastic spatial simulations. However, our findings indicate that the stQSSA can introduce substantial errors in heterogeneous environments typical of spatial SSA. This error arises because the stQSSA may satisfy the validity conditions globally in the SSA, but not within the confined regions where DNA is localized. In contrast, the slQSSA, derived by analytically solving the CME under the assumption of a small number of DNA binding sites, consistently provides accurate estimates. Therefore, utilizing the slQSSA is recommended to reduce simulation time in spatiotemporal stochastic models.

## 4. Materials and Methods

### 4.1. Mathematical model describing gene regulatory networks

#### 4.1.1. One binding site gene regulatory network model

To describe the simplest case of a gene regulatory network (Fig. 1A), we used a model with one binding site on DNA (*D*_*X*_, *X ∈* {0, 1}):

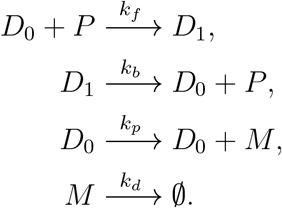

DNA with a single binding site reversibly binds to a transcriptional repressor (*P*), characterized by binding and unbinding rates *k*_*f*_ and *k*_*b*_, respectively. Specifically, unoccupied DNA (*D*_0_) can bind to *P* to form occupied DNA (*D*_1_). mRNA (*M*) is transcribed only from *D*_0_ at a rate of *k*_*p*_. *M* decays at a rate of *k*_*d*_.

#### 4.1.2. Two binding site gene regulatory network model and its simplification

DNA with multiple binding sites exhibits more complex gene regulation compared to DNA with a single binding site [27, 63, 64]. To investigate these properties, we utilized the following model that describes the binding of *P* to DNA with two binding sites:

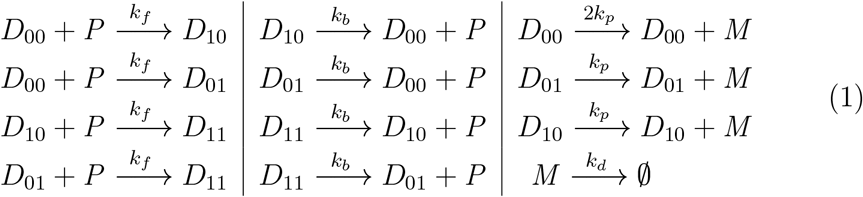

In this model, DNA with two binding sites reversibly binds to *P* with binding rate *k*_*f*_ and unbinding rate *k*_*b*_. Unoccupied DNA (*D*_00_) can bind *P* at either site, forming *D*_01_ or *D*_10_. These singly-occupied states can then bind another *P* at the remaining site, forming fully occupied DNA (*D*_11_). *M* transcription occurs at a rate of 2*k*_*p*_ from *D*_00_, at a rate of *k*_*p*_ from *D*_01_ or *D*_10_, and not at all from *D*_11_. This mechanism can be described by the following chemical master equations (CMEs) [27, 65]:

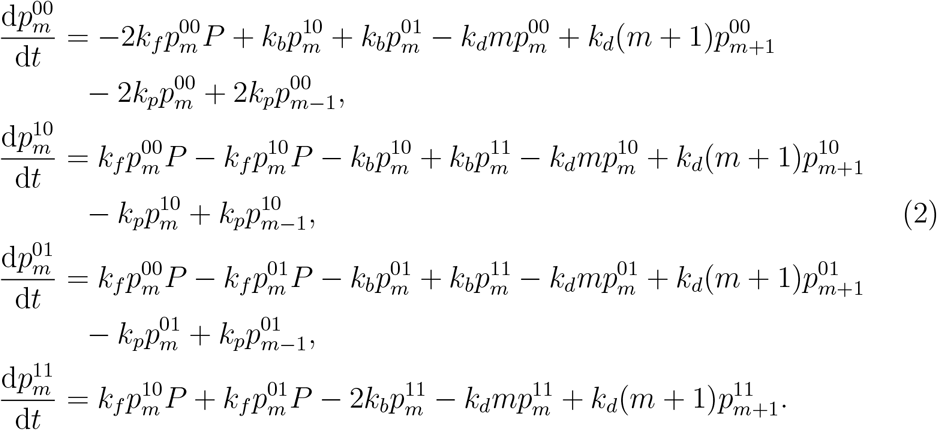

Here, 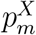 represents the joint probability that DNA is in state *D*_*X*_ with *m* mRNAs, *X ∈* {00, 01, 10, 11} and *P* denotes the number of repressors. To represent the binding and unbinding of *D*_*X*_ with *P*, we respectively used 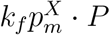 and 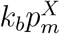 that are the stochastic transitions between DNA states. Transcription is activated in the remaining DNA state at a rate *k*_*p*_. For example, 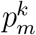 decrease at the rate 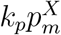 and increases at the rate 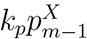. However, when both binding sites are occupied (i.e., *D*_11_), transcription is fully repressed, meaning that mRNA transcription has no impact on the change in 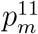. To represent the degradation of each mRNA, 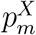 decreases at the rate 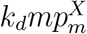 and increases at the rate 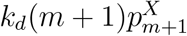.

By substituting 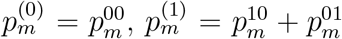, and 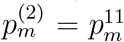, we can simplify these equations (Eq. 2) into the following compact form:

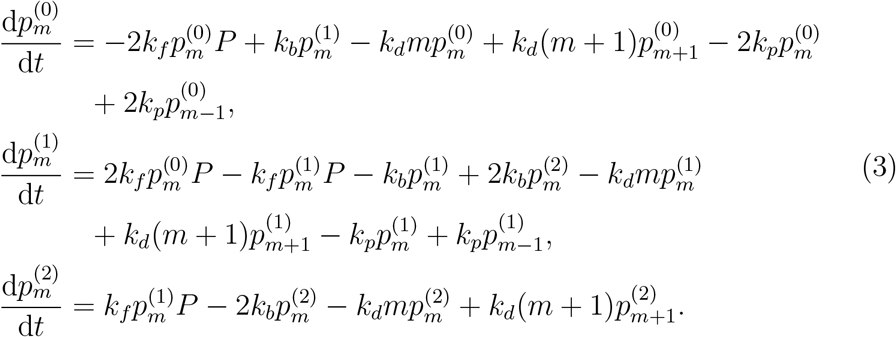

Note that Eq. 3 is identical to the following simplified system where the number of unoccupied binding sites of DNA (*D*) = 2, where we define 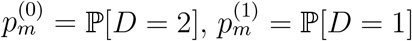 and 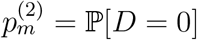 :

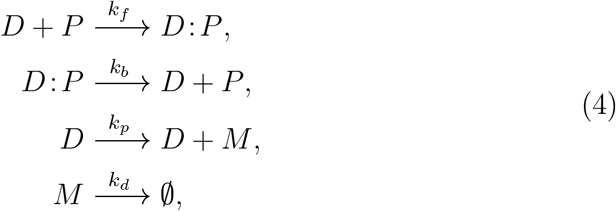

where *D* : *P* represents the complex form of *D* and *P*. Instead of simulating the complicated model, we implemented the equivalent simplified model (Eq. 4).

### 4.2. Stochastic simulation of heterogeneous environments using compartment-based Gillespie algorithm

In order to model the gene regulation (Eq. 4) in heterogeneous environments, we employed a spatial stochastic simulation algorithm, specifically using a compartment-based Gillespie algorithm [31]. To realize this algorithm, we first partitioned the one-dimensional system into *K* compartments of size *h* = *L/K*. We then simulated the reaction in each compartment and the diffusion between these compartments. Across compartments, repressors can diffuse at a rate of *d* = *δ/h*^2^, where *δ* is the diffusion coefficient of the repressor, while DNA cannot diffuse. Additionally, the diffusion of the repressor can be described by the following linear reactions: 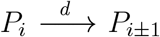, where *P*_*i*_ denotes the number of repressors in the *i −*th compartment. Within each compartment, reactions are described by Eq. 4. In other words, repressors can bind to binding sites of DNA only when they are in the same compartment. Thus, the propensity functions of the stochastic reaction-diffusion equation can be obtained as follows [14, 31]:

**Table 1:** Propensity functions of the full model describing the gene regulatory network used in the spatial SSA.

| Propensity functions |  | Propensity |
| --- | --- | --- |
| $D_i + P_i \xrightarrow{k_f} D:P_i$ | $i = 1, \dots, K$ | $\frac{k_f}{\Omega} D_i P_i$ |
| $D:P_i \xrightarrow{k_b} D_i + P_i$ | $i = 1, \dots, K$ | $k_b D:P_i$ |
| $D_i \xrightarrow{k_p} D_i + M_i$ | $i = 1, \dots, K$ | $k_p D_i$ |
| $M_i \xrightarrow{k_d} \emptyset$ | $i = 1, \dots, K$ | $k_d M_i$ |
| $P_i \xrightarrow{d} P_{i-1}$ | $i = 2, \dots, K$ | $d P_i$ |
| $P_i \xrightarrow{d} P_{i+1}$ | $i = 1, \dots, K-1$ | $d P_i$ |

For the spatial SSA of Figs. 2 and 3, we set *K* to 30, resulting in the compartment size *h* = 0.33*µm*. Here, we set the size of the nucleus to *L* = 10*µm*, which is the typical size of a mammalian nucleus [59]. The volume of each compartment (Ω) was set to 1, resulting in the size of the whole system becoming Ω*×K* = 30. At the boundary compartments (*i* = 1 and *i* = *K*), diffusion is constrained: no leftward diffusion and rightward diffusion occur at *i* = 1 and *i* = *K*, respectively. In Fig 2, to ensure tight binding between DNA and *P*, we set the binding and unbinding rates as *k*_*f*_ */*Ω = 5000*/*Ω*s*^*−*1^, *k*_*b*_ = 100*s*^*−*1^, respectively (resulting in a dissociation constant 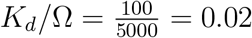 [14]. Additionally, *k*_*p*_ = 50*s*^*−*1^, *k*_*d*_ = 0.001*s*^*−*1^. The simulation was conducted for *T*_*max*_ = 50*s*.

We set *D*_*i*_(0) and *P*_*i*_(0) at *t* = 0 as follows: In Fig. 2A-D, we used initial conditions where DNA with one binding site is immobile in the center of the nucleus and *P* has a uniform distribution in the nucleus: *D*_15_(0) = 1, *D*_*i*_(0) = 0 for *i*≠ 15, and *P*_*i*_(0) = 1 for *i* = 1 … *K*. In Fig. 2A-B, we set *P* to diffuse quickly between compartments at a rate of *d* = 180.4*s*^*−*1^, which corresponds to the diffusion coefficient of *δ* = 20 *µm*^2^*/s*, which falls within the typical range for protein diffusion coefficients [66]. However, in Fig. 2C-H and Fig. 3C-D, we slowed down the diffusion rate to *d* = 1.8*s*^*−*1^, which corresponds to the diffusion coefficient of PER2 protein, *δ* = 0.2*µm*^2^*/s* [67]. In Figs. 2E and 3C, we only changed the initial distribution of *P* to have a nonuniform distribution in the nucleus from Fig. 2C: *P*_*i*_(0) = 1 for *i* = 11, …, 20, and *P*_*i*_(0) = 5 for *i* = 11, …, 20. For Figs. 2G and 3D, we only changed the initial conditions for *D*, without changing the initial condition for *P* from Fig. 2C: *D* with two binding sites is immobile in the center of the nucleus as *D*_15_(0) = 2, *D*_*i*_(0) = 0 for *i*≠ 15.

### 4.3. Stochastic simulation of homogeneous environments using Gillespie algorithm

To simulate gene regulation in homogeneous environments, we utilized the stochastic simulation algorithm (SSA), which is also known as the Gillespie algorithm [68]. Different from the spatial SSA, the SSA ignores spatial heterogeneity. For example, *P* can bind to *D* at any location within the nucleus. In contrast, the spatial SSA model restricts *P* to binding with *D* only when they are sufficiently close to each other.

Thus, we calculated the propensity functions of the full model based on the reactions as follows: For the SSA simulation, Ω was set to 30 to match the volume of the whole system size in the spatial SSA. And for Fig. 2, we utilized the same parameters as the previous section. For Fig. 2, the following initial conditions were used: [*D*(0), *P* (0)] = [1, 30] for Fig. 2A and C, [*D*(0), *P* (0)] = [1, 110] for Fig. 2E, and [*D*(0), *P* (0)] = [2, 30] for Fig. 2G.

**Table 2:** Propensity functions of the full model describing the gene regulatory network used in the SSA.

| Reaction | Propensity |
| --- | --- |
| $D + P \xrightarrow{k_f} D:P$ | $\frac{k_f}{\Omega} DP$ |
| $D:P \xrightarrow{k_b} D + P$ | $k_b D:P$ |
| $D \xrightarrow{k_p} D + M$ | $k_p D$ |
| $M \xrightarrow{k_d} \emptyset$ | $k_d M$ |

### 4.4. Derivation of two reduction model equations: slQSSA and stQSSA

Although the spatial SSA is required to describe gene regulation with spatial heterogeneity, it is significantly more computationally expensive than the traditional SSA. To address this issue, we applied the QSSA, which approximates the amount of free DNA that rapidly reaches a quasi-steady state (QSS) due to the fast reaction, thereby allowing us to focus only on the slow reactions. To apply QSSA to stochastic systems, it is necessary to solve the CMEs to obtain the average number of stationary states. While this is typically challenging, it can be computed analytically when the number of binding sites is low. This approach is known as the slQSSA [14]. In our study, we applied the slQSSA within the spatial SSA framework, considering three states based on the number of occupied binding sites as follows:

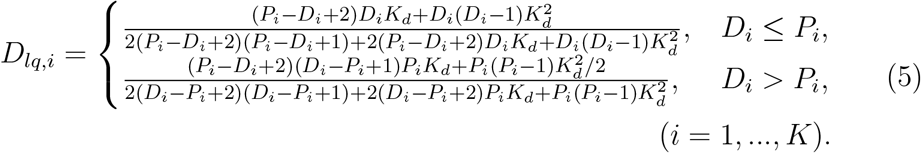

As the number of binding sites increases, the slQSSA equation (Eq. 5) becomes more complex, making it more difficult to interpret. As an alternative, previous studies have used the stQSSA [10, 11, 12, 13, 14, 42, 44, 45, 46, 47, 48, 49, 50, 51]. The stQSSA *D*_*tq,i*_ can be derived using the concentration-based free DNA obtained from the deterministic tQSSA:

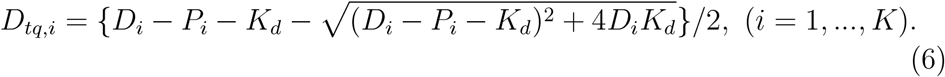

While it has been believed that the stQSSA accurately captures stochastic dynamics at a low computational cost [10, 11, 12, 13, 14, 42, 44, 45, 46, 47, 48, 49, 50, 51], recent research showed that stQSSA can be inaccurate when two species tightly bind and their molar ratio is *∼*1:1 [14].

Based on the obtained *D*_*lq,i*_ and *D*_*tq,i*_, we calculated the propensity functions to verify that the two reduced models accurately approximate the full model. Here, *P*_*lq,i*_ and *P*_*tq,i*_ represent the number of the free repressor molecules in the *i−*th compartment by slQSSA and stQSSA, respectively. Since bound repressors cannot diffuse, the free repressor should be predicted by using slQSSA or stQSSA. Specifically, the number of free repressor molecules is calculated by subtracting the number of bound *P* (*D* : *P*_*lq,i*_ or *D* : *P*_*tq,i*_) from the total *P* obtained from the simulation (*P*_*i*_), using *P*_*lq,i*_ = *P*_*i*_ *− D* : *P*_*lq,i*_, *P*_*tq,i*_ = *P*_*i*_ *− D* : *P*_*tq,i*_, where *D* : *P*_*lq,i*_ = *D*_*i*_ *− D*_*lq,i*_, *D* : *P*_*tq,i*_ = *D*_*i*_ *− D*_*tq,i*_.

**Table 3:** Propensity functions of the reduced (stQSSA and slQSSA) models describing the gene regulatory network used in the spatial SSA.

| Reaction | Propensity functions |  |
| --- | --- | --- |
|  | slQSSA | stQSSA |
| $D_i \xrightarrow{k_p} D_i + M_i, \quad i = 1, \dots, K$ | $k_p D_{lq,i}$ | $k_p D_{tq,i}$ |
| $M_i \xrightarrow{k_d} \emptyset, \quad i = 1, \dots, K$ | $k_d M_i$ | |
| $P_i \xrightarrow{d} P_{i-1}, \quad i = 2, \dots, K$ | $d P_{lq,i}$ | $d P_{tq,i}$ |
| $P_i \xrightarrow{d} P_{i+1}, \quad i = 1, \dots, K-1$ | $d P_{lq,i}$ | $d P_{tq,i}$ |

To simulate two reduction models (slQSSA and stQSSA) in Fig. 3, we set *K* to 30 and divided the 1-dimensional domain [0, 10] into *K* = 30 compartments of length *h* = 0.03*µm*. We used Ω = 1, *k*_*f*_ */*Ω = 5000*/*Ω*s*^*−*1^, *k*_*b*_ = 100*s*^*−*1^, *k*_*p*_ = 50*s*^*−*1^, and *k*_*d*_ = 0.001*s*^*−*1^. The diffusion rate was set to *d* = 0.01*s*^*−*1^, and the simulation ran for *T*_*max*_ = 50*s*. For Fig. 3C-D, *P* was set to diffuse slowly between compartments at a rate of *d* = 0.018*s*^*−*1^. In Fig. 3C-D, the QSSA models use the same initial conditions as the full model.

### 4.5. Stochastic simulation of molecular oscillation

We constructed a gene regulatory model with a negative feedback loop to investigate the periodicity of biological oscillations across three models [69, 70, 71, 72, 73]: the full, the slQSSA, and the stQSSA. Unlike the previous model, this domain represents the entire cell. Specifically, the cell is 30*µm* in size, with a 10*µm* region at the center representing the nucleus, which is the typical size of mammalian cells [59]. We assume that the DNA is immobile within the nucleus, while repressors can diffuse in and out of the nucleus. The propensity functions for the oscillation model are as follows:

For Fig. 4, we used *K* = 30 and thus *h* = 1*µm*, and we assumed that *d* = 0.2*s*^*−*1^ as in Fig. 2C-H. We set Ω = 1, and the rate constants were set to *k*_*f*_ */*Ω = 10000*/*Ω*s*^*−*1^, *k*_*b*_ = 100*s*^*−*1^ to ensure tight binding (*K*_*d*_ = 0.02), and *k*_*p*_ = 10*s*^*−*1^, *k*_*d*_ = 0.01*s*^*−*1^. he simulation ran for a total duration of *T*_*max*_ = 5000*s*. For the initial conditions, DNA with two binding sites was assumed to be immobile in the center of the nucleus, and repressors had a uniform distribution in the nucleus: *D*_15_(0) = 2, *D*_*i*_(0) = 0 for *i≠15*, and *P*_*i*_(0) = 1 for *i* = 1 … *K*.

**Table 4:** Propensity functions of the full and reduced (stQSSA and slQSSA) models describing biological oscillator used in the spatial SSA.

| Reaction | Propensity functions |  |  |
| --- | --- | --- | --- |
|  | Full | slQSSA | stQSSA |
| $D_i + P_i \xrightarrow{k_f} D:P_i \quad i = 1, \dots, K$ | $\frac{k_f}{\Omega} D_i P_i$ | - | - |
| $D:P_i \xrightarrow{k_b} D_i + P_i \quad i = 1, \dots, K$ | $k_b D:P_i$ | - | - |
| $D_i \xrightarrow{k_p} D_i + P_i \quad i = 1, \dots, K$ | $k_p D_i$ | $k_p D_{lq,i}$ | $k_p D_{tq,i}$ |
| $P_i \xrightarrow{k_d} \emptyset \quad i = 1, \dots, K$ | $k_d P_i$ | | |
| $D:P_i \xrightarrow{k_d} D_i \quad i = 1, \dots, K$ | $k_d D:P_i$ | | |
| $P_i \xrightarrow{d} P_{i-1} \quad i = 2, \dots, K$ | $d P_i$ | $d P_{lq,i}$ | $d P_{tq,i}$ |
| $P_i \xrightarrow{d} P_{i+1} \quad i = 1, \dots, K - 1$ | $d P_i$ | $d P_{lq,i}$ | $d P_{tq,i}$ |

### 4.6. Quantifying the period of molecular oscillations

In Fig. 4D, we quantified the period of oscillation of *P* by utilizing the decaying cosine function following previous studies [57, 28, 58]. First, we calculated the autocorrelation function *C*(*s*) of the total protein concentration in the cell over time, *P* (*t*), using the following formula: 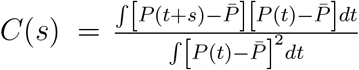, where 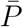 is the time average of *P* (*t*). We then fitted this to a decaying cosine function 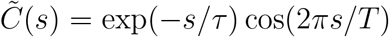 to estimate the period of oscillation *T*, where the correlation time *τ* describes how fast the *C*(*s*) exponentially decays. To ensure accuracy, the period was calculated using the simulation data from the last 2,500*s* of a 5,000*s* run, with the first 2,500*s* excluded to eliminate the influence of transient dynamics. To obtain the period distribution in Fig. 4D, we independently calculated the period of 1,000 iterative simulations.

## Acknowledgments

This work was supported by the Institute for Basic Science (grant no. IBS-R029-C3 to JKK), the National Research Foundation of Korea (NRF) grant funded by the Korea government (MSIP) (grant no. RS-2024-00342949 to SL), and Regional Innovation Strategy (RIS) through the NRF funded by the Ministry of Education (MOE) (grant no. 2021RIS-004 to SL). The funders had no role in study design, data collection and analysis, decision to publish, or preparation of the manuscript.

## Author Contributions

Conceptualization: Seunggyu Lee, Jae Kyoung Kim

Data Curation: Seolah Shin, Seok Joo Chae

Formal analysis: Seolah Shin, Seok Joo Chae, Seunggyu Lee, Jae Kyoung Kim

Funding acquisition: Seunggyu Lee, Jae Kyoung Kim.

Investigation: Seolah Shin, Seok Joo Chae, Seunggyu Lee, Jae Kyoung Kim Methodology: Seolah Shin, Seok Joo Chae, Seunggyu Lee, Jae Kyoung Kim Project administration: Seunggyu Lee, Jae Kyoung Kim.

Resources: Seunggyu Lee, Jae Kyoung Kim

Software: Seolah Shin, Seok Joo Chae

Supervision: Seunggyu Lee, Jae Kyoung Kim

Validation: Seolah Shin, Seok Joo Chae, Seunggyu Lee, Jae Kyoung Kim Visualization: Seolah Shin, Seok Joo Chae, Seunggyu Lee, Jae Kyoung Kim

Writing – original draft: Seolah Shin, Seok Joo Chae, Seunggyu Lee, Jae

Kyoung Kim

Writing – review & editing: Seolah Shin, Seok Joo Chae, Seunggyu Lee, Jae

Kyoung Kim

